# Inflammation modulates regeneration in the acute or chronically damaged zebrafish retina

**DOI:** 10.1101/2021.10.06.463331

**Authors:** Maria Iribarne, David R. Hyde

## Abstract

Unlike mammals, zebrafish regenerate in response to retinal damage. Because microglia are activated by retinal damage, we investigated their role during regeneration following acute or chronic damage. At three weeks-post-fertilization (wpf), fish exhibiting NMDA-induced acute damage or cone photoreceptor-specific chronic degeneration, the *gold rush (gosh)* mutant, displayed reactive microglia and Müller glia proliferation. Retinas treated to inhibit the immune response lacked reactive microglia and possessed fewer PCNA-positive cells, while LPS treatment increased microglia and PCNA-labeled cells. NMDA-injured retinas upregulated the expression of il-1β and tnf-α pro-inflammatory cytokine genes, followed by increased expression of il-10 and arg1 anti-inflammatory/remodeling cytokine genes. An early and transiently TNF-α pro-inflammatory microglia population was identified in the NMDA-damaged retina. In contrast, *gosh* mutant retinas exhibited a mild increase of pro-inflammatory cytokine gene expression concurrently with a greater increased in anti-inflammatory/remodeling cytokine gene expression. Few TNF-α pro-inflammatory microglia were observed in the *gosh* retina. How inflammation regulates regeneration in zebrafish would provide important clues towards improving the therapeutic strategies for repairing injured mammalian tissues.

## Introduction

Most vertebrates, including humans, can not to regenerate retinal neurons that are lost due to traumatic injury or degenerative disease. In contrast, lower vertebrates, such as zebrafish, possess an extraordinary regenerative ability, which restores the normal function to the damaged retina (Gemberling, Bailey, Hyde, & Poss, 2013; Hoang et al., 2020; Iribarne, 2019). Upon neuronal loss in the zebrafish retina, Müller glia reprogram to a retinal progenitor cell-like state and re-enter the cell cycle to generate neuronal progenitor cells (NPCs). These NPCs continue to proliferate and migrate to the site of neuronal damage and differentiate into the neuronal types that were lost (Bernardos, Barthel, Meyers, & Raymond, 2007; Fausett & Goldman, 2006; Fimbel, Montgomery, Burket, & Hyde, 2007; Yurco & Cameron, 2005). Recent studies in the mouse retina were able to induce a small proliferative response by Müller glia after either the overexpression or loss of certain transcription factors, which were identified in the regenerative response in the zebrafish retina (Elsaeidi et al., 2018; Jorstad et al., 2017; Yao et al., 2018). Thus, understanding mechanisms by which zebrafish can regenerate injured tissue may provide strategies for stimulating mammalian retina regeneration.

Recent evidence suggests that the innate immune system can modulate the regenerative response following zebrafish neuronal damage. Acute and transient inflammation is necessary to induce a regenerative response in the adult zebrafish telencephalon (Kyritsis et al., 2012). Depleting or inhibiting microglia functions following rod photoreceptor ablation in zebrafish larvae blocked the regenerative response by Müller glia (White et al., 2017). Pharmacological treatment with either dexamethasone or PLX3397 reduced the number of proliferating progenitor cells in adult zebrafish following various retinal injuries (Conedera, Pousa, Mercader, Tschopp, & Enzmann, 2019; Silva et al., 2020; Zhang et al., 2020). Although the involvement of inflammation during retinal regeneration has been reported, its molecular mechanism of modulating Müller glia proliferation remains elusive.

Inflammation is a dynamic process involving the recruitment of inflammatory cells and the secretion of pro-inflammatory cytokines and molecular mediators. The resolution of the inflammation is critical to avoid tissue damage (Bosak, Murata, Bludau, & Brand, 2018; Iribarne, 2021). M1-like macrophages are pro-inflammatory cells associated with the first phases of inflammation and typically express IL-1β and TNF-α; while M2-like macrophages are involved in the resolution of inflammation and tissue remodeling response and express IL-10 and TGF-β1 (Nguyen-Chi et al., 2015). The differential expression of cytokines and chemokines, as well as the expression of receptors, defines the polarization state of macrophages. Several studies in zebrafish suggest that the activation and duration of pro-inflammatory signals and the subsequent resolution are critical in creating an instructive microenvironment for tissue regeneration. For instance, a transient inflammatory response mediated by IL-1β is required for proper regeneration of the zebrafish fin fold, where macrophages are responsible for the attenuation of IL-1β expression (Hasegawa et al., 2017). Similarly, an interplay between IL-1β and TNF-α from neutrophils and macrophages is necessary for the regeneration of the injured spinal cord in zebrafish larvae (Tsarouchas et al., 2018). However, it remains unknown what cytokine profiles are expressed during the regeneration process and whether microglia switch from pro-inflammatory to resolution state in the damaged retina.

Most previous retinal regeneration studies have employed acute damage in the adult retina, such as strong light exposure (Bernardos et al., 2007; Vihtelic & Hyde, 2000), retinal puncture (Fausett & Goldman, 2006), chemical ablation (Fimbel et al., 2007; Powell, Cornblath, Elsaeidi, Wan, & Goldman, 2016), or ectopic expression of a toxic transgene, such as nitroreductase (Hagerman et al., 2016; Montgomery, Parsons, & Hyde, 2010). Acute damage leads to rapid loss of retinal cells that resemble traumatic injury in human patients. On the other hand, retinal regeneration studies using chronic damage models in the zebrafish retinas are limited (Iribarne, Hyde, & Masai, 2019; Iribarne et al., 2017; Morris, Scholz, Brockerhoff, & Fadool, 2008; Nishiwaki et al., 2008; Sherpa, Hunter, Frey, Robison, & Stenkamp, 2011; Turkalj et al., 2021). These chronic degeneration mutants better model human genetic diseases, which usually begin during embryogenesis or childhood and take a long time to completely degenerate. Notably, both acute and chronic retinal damages in zebrafish can induce a Müller glia-dependent regenerative response. While several acute damage studies have been performed focusing on the role of inflammation during the regeneration process, similar studies using chronic retinal models have not been reported. Therefore, we examined the regenerative response following acute and chronic retinal damage with a focus on the role of inflammation. Understanding how inflammation regulates regeneration in either acute or chronic retinal damage in zebrafish would provide important insights to improve the therapeutic strategies for repairing injured mammalian tissues that do not have an inherent regenerative capacity.

## Results

### Damaged retinas stimulate microglial activation and Müller glia proliferation

We took advantage of two different neuronal injury paradigms to investigate whether inflammation regulates Müller glia proliferation. We adapted a NMDA-mediated neurotoxicity model that selectively damages neurons in the adult zebrafish inner nuclear layer (INL) and ganglion cell layer (GCL), but spares photoreceptors to generate an acute injury (Luo et al., 2019; Powell et al., 2016). The genetic mutant *gosh*, which exhibits a progressive cone photoreceptor degeneration and regeneration, was used as a chronic retinal damage model (Iribarne et al., 2019; Iribarne et al., 2017).

Notably, larval zebrafish possess only the innate immune system, which allows for the study of innate responses in isolation. In contrast, the adaptive immune system develops around 4 to 6 wpf (Yoder, Nielsen, Amemiya, & Litman, 2002). Thus, we used 3 wpf fish to study the innate immune system response in the two retinal injury models. We used the transgenic line *Tg(mpeg1:GFP)*, which expresses GFP specifically in macrophages, to monitor the macrophages/microglia activation response (Ellett, Pase, Hayman, Andrianopoulos, & Lieschke, 2011). Control retinas at 3 wpf showed GFP-positive macrophages/microglia in different retinal layers and displayed ramified morphology with long processes (Figure 1A, arrows). At 72 hours following intravitreal NMDA injection, there were an increased number of GFP-expressing cells, primarily in the injured inner retina, which switched to an ameboid cell shape (Figure 1B, G; control retinas: 5.43 ± 0.54 GFP+ cells; NMDA: 44.54 ± 1.98; p<0.001). To distinguish between microglia or infiltrated macrophages, we stained with the 4C4 monoclonal antibody that specifically labels microglia, but not peripheral macrophages (Suppl. Figure 1). Control retinas showed only one population of microglia, while NMDA-induced microglia increased, accompanied by a few recruited peripheral macrophages in the retina.

**Figure 1.**
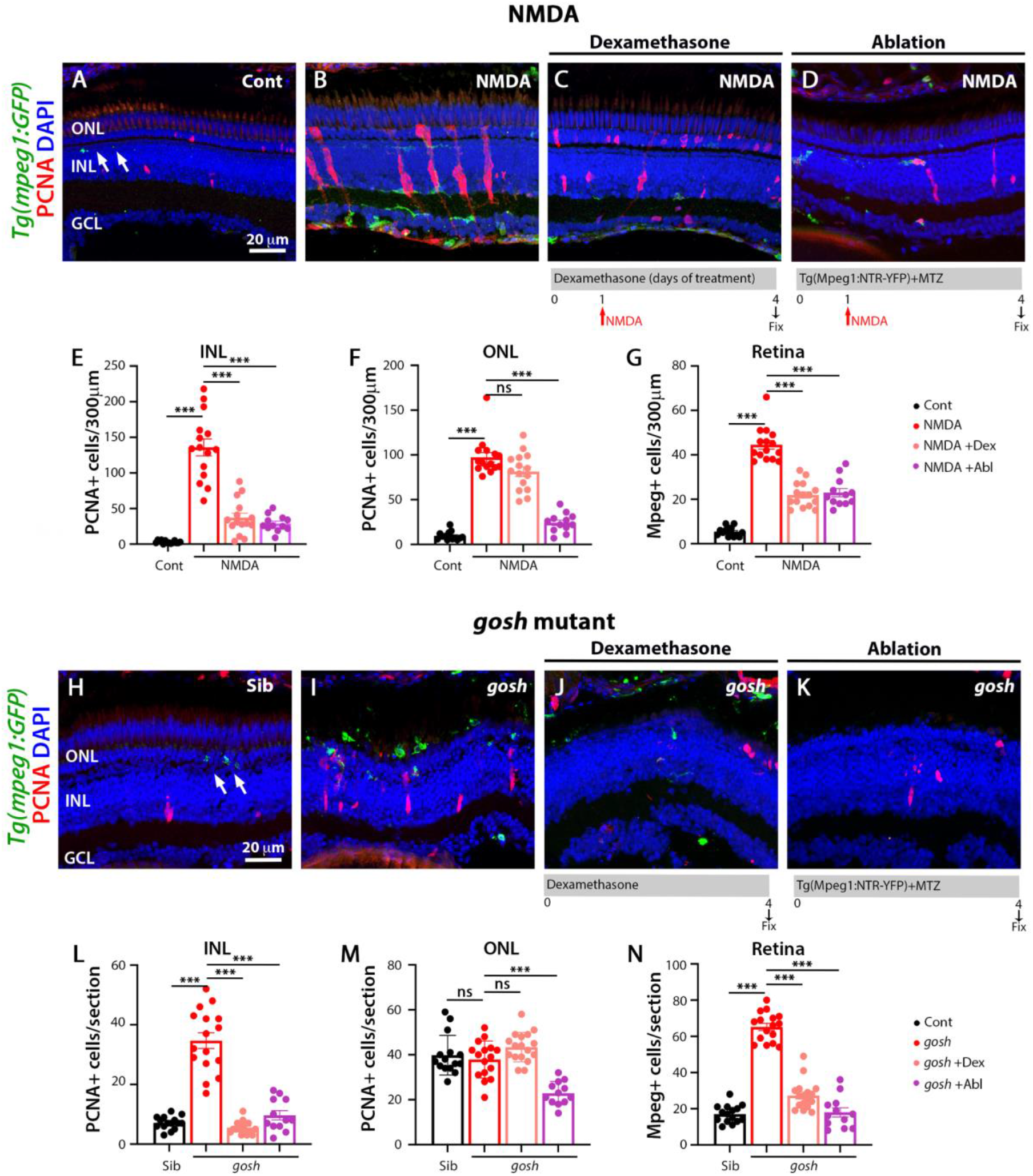
Suppressing microglia affect Müller glia proliferation in NMDA-injured retina and *gosh* mutants. Labeling of 3-wpf fish with NMDA-injured retina after 72 hours and *gosh* mutant retinas with anti-PCNA antibody. Microglia are visualized with the *Tg(mpeg1:GFP)* transgenic line, and nuclei are counterstained with DAPI. Wild-type retinas display few thin and ramified microglia through the retina (arrows), and PCNA-positive cells are observed in the ONL and INL, corresponding to rod precursors and Müller glia/NPCs, respectively (A, E-G). In the NMDA-injured retina, microglia are in high numbers in the inner part of the retina, and PCNA expression is strongly stimulated (B). Dexamethasone treatment (C) or microglia ablation (D) in the NMDA-damaged retinas reduces the number of microglia and PCNA expressing cells in INL and ONL. Wild-type sibling retinas display some ramified microglia (arrows) and few PCNA-positive cells (H). *gosh* mutant retinas have activated microglia located in the ONL (I). PCNA expressing cells are present in the INL and ONL. Dexamethasone treatment (J) or macrophages ablation (K) reduces PCNA expressing cells in INL *gosh* retinas. Histograms display the quantification of the number of PCNA-labeled cells (E-F) or GFP-positive macrophages (G) in 300 m of the central region of the retina for NMDA or in the entire section of the retina for *gosh* mutant (L-N). Bars and lines indicate mean ± SEM, n: 12-17. Black bars: wild-type sibling; red bars: NMDA-injected retinas or *gosh* mutant; pink bars: dexamethasone treatment; purple bars: ablated macrophages. One-way ANOVA with Tukey’s multiple comparisons test was applied for all the graphs (ns p>0.05; ***p<0.001).

We also co-labeled wild-type and NMDA-damaged retinas with anti-PCNA antibody, a cell proliferation marker. In the wild-type retina, a small number of PCNA-positive cells were observed in the INL and outer nuclear layer (ONL), which likely correspond to Müller glia and rod progenitor cells, respectively (Figure 1A). These proliferating cells are the source of persistent neurogenesis, where Müller glia divide asymmetrically and infrequently to produce rod progenitor cells, which migrate to the ONL and are committed to differentiate into rod photoreceptors (Lahne, Li, Marton, & Hyde, 2015; Nagashima, Barthel, & Raymond, 2013). Upon NMDA injection, PCNA-positive cell numbers increased in the INL and ONL (Figure 1A, B, E, F; control retinas, INL: 3.07 ± 0.52 and ONL: 9.50 ± 1.25; NMDA INL: 136.00 ± 11.88, p<0.001; and ONL: 97.40± 5.39, p<0.001). Clusters of PCNA-positive cells, which are formed by Müller glia and the Müller glia-derived neuronal progenitor cells (NPCs) arranged in a vertical column, could often be observed. To corroborate that these PCNA-positive INL cells correspond to Müller glia, we used the *Tg(gfap:GFP)* line to visualize Müller glia and stained for PCNA at 48 hours post injury (hpi) (Suppl. Figure 2). Control retinas exhibited low levels of PCNA-positive Müller glia, which correspond to persistent neurogenesis. NMDA-injured retinas showed several Müller glia stained with PCNA to induce a regenerative response.

Previously, we showed that the *gosh* mutant underwent photoreceptor degeneration (Iribarne et al., 2017). *gosh* mutant retinas at 3 wpf displayed a very thin photoreceptor layer, where cones form a discontinuous layer, and the central retina being the worst affected. In our previous characterization of the regeneration process in the *gosh* mutant, we found that Müller glia did not proliferate at 3 wpf, but were proliferating by 5 wpf (Iribarne et al., 2019). In this study, we fed the zebrafish larvae with rotifers from 4 dpf to 12 dpf; under these conditions, the larvae overcame the proliferation delay that we observed previously, with the *gosh* mutant possessing a proliferative response at 3 wpf. In wild-type sibling retinas, microglia were detected mainly in the inner plexiform layer but also frequently in the ONL (Figure 1H). In *gosh* mutant retinas, microglia displayed a stronger GFP intensity, a greater number of cells, and were localized primarily in the ONL, where photoreceptors were dying, and in the outer segment region (Figure 1I, N; sibling wild-type retinas: 17.07 ± 1.26 GFP+ cells; *gosh* mutant: 65 ± 2.00; p<0.001). The 4C4 antibody staining in the transgenic Tg(mpeg1:GFP) fish demonstrated that sibling retinas contained only microglia, while *gosh* mutant retinas possessed mostly microglia and a few peripheral macrophages that infiltrated into the retina (Suppl. Figure 3).

PCNA immunostaining revealed cells in the INL and ONL that formed small clusters (Figure 1H, I, L, M; sibling retina INL: 7.13 ± 0.56 and ONL: 39.8± 2.28 PCNA+ cells; *gosh* mutant INL: 34.69 ± 2.63, p<0.001; and ONL: 37.88 ± 2.05, p=0.89). The *Tg(gfap:GFP)* line was introduced into the *gosh* background to assess if Müller glia were dividing (Suppl. Figure 4). Again, wild-type sibling retinas displayed a few PCNA-positive cells in the INL and ONL. *gosh* mutant retinas showed a greater number of PCNA-labeled cells, some of which were in the INL and co-labeled with GFP, indicating that those cells were Müller glia re-entering the cell cycle. Taking together these data revealed that both the acute and chronic retinal damage models induced activation of inflammatory cells and a regenerative response involving Müller glia.

### Dexamethasone or nitroreductase-metronidazole ablation treatment reduces microglial activation and Müller glia proliferation in injured retinas

To evaluate whether microglia play a role in the regenerative response following either acute or chronic retinal damage, we used two independent methodologies to suppress microglia in the retina: 1) the anti-inflammatory glucocorticoid dexamethasone (Dex) to inhibit microglia/macrophages (Silva et al., 2020; White et al., 2017) and 2) the nitroreductase-metronidazole system to ablated microglia/macrophages (Petrie, Strand, Yang, Rabinowitz, & Moon, 2014). The dexamethasone treatment started one day before the NMDA injection and continued for four days. NMDA retinas treated with dexamethasone had a reduced number of microglia compared with the NMDA group (Figure 1C, G; NMDA+Dex: 21.93 ± 1.45, p<0.001). These retinas also possessed a reduced number of PCNA-positive cells in the INL, but similar levels of PCNA-positive cells in the ONL relative to NMDA retinas (Figure 1C, E, F; PCNA+ NMDA Dex INL: 37.20 ± 6.31, p<0.001 and ONL: 81.33 ± 5.43, p=0.06). We ablated macrophages employing the *Tg(mpeg1:NTR-YFP)* line that expresses the nitroreductase enzyme in macrophages, which, when treated with the pro-drug metronidazole (MTZ) leads to the apoptosis of these cells. MTZ treatment started one day before the NMDA injection and continued for four days. NMDA retinas treated with MTZ displayed fewer macrophages/microglia in the retina than the NMDA group (Figure 1D, G; NMDA ablation: 23 ± 1.84, p<0.001). The number of PCNA-positive cells was also reduced in these retinas, both in the INL and ONL compared to NMDA retinas (Figure 1D-F; NMDA ablation INL: 29.08 ± 3.29, p<0.001 and ONL: 24.25 ± 3.05 p<0.001).

In *gosh* mutant retinas treated four days with dexamethasone, microglia cells appeared thinner, ramified, and less numerous than *gosh* retinas (Figure 1J, N; *gosh* mutant+Dex: 27.2 ± 1.94 mpeg+ cells; p<0.001) compared to *gosh* mutants. Microglia did not accumulate in the ONL or outer segment layer (Figure 1J). *gosh* mutant retinas treated with dexamethasone showed very few PCNA-labeled cells in the INL, but maintained a high number in the ONL (Figure 1J, L, M; *gosh* mutant+Dex INL: 5.53 ± 0.50, p<0.001 and ONL: 43.35 ± 1.59, p=0.17). The *gosh* mutant carrying on the *Tg(mpeg1:NTR-YFP)* transgene were treated four days with the pro-drug MTZ. Microglia ablation efficiently reduced the number of microglia in the *gosh* mutant (Figure 1K, N: *gosh* mutant ablation: 18.5 ± 2.8, p<0.001) compared to *gosh* mutants. Microglia ablated in *gosh* mutant retinas exhibited few PCNA-positive cells the ONL (Figure 1K, L, M; *gosh* mutant ablation INL: 11.08 ± 2.3 PCNA+ cells and ONL: 22.83 ± 1.48). All these data suggest that either dexamethasone treatment or nitroreductase-MTZ ablation system can effectively reduce the number of microglia and the number of Müller glia re-entering the cell cycle in acute or chronic retinal damage.

### LPS increases the number of microglia and proliferating Müller glia in wild-type and injured retinas

To induce inflammation, bacterial lipopolysaccharide (LPS) was injected in the acute and chronic retinal injury models. Subsequently, microglia/macrophage activation and Müller glia proliferation were evaluated in cryosections. LPS-treated retinas showed an increased number of microglia presence compared to PBS control retinas that was not statistically significant (Figure 2A, B, H; control retinas: 5.43 ± 0.53 mpeg1+ cells; control retinas+LPS: 11.79 ± 0.99, p=0.69). NMDA-induced damage showed an increased number of microglia after LPS treatment (Figure 2C, D, H; NMDA retinas: 43.07 ± 1.55 mpeg+ cells; NMDA retinas+LPS: 61.53 ± 2.14, p<0.001). In addition, LPS injection led to an increase of number of proliferating Müller glia and NPCs in the NMDA damaged retinas compared to PBS-treated retinas (Figure 2B, D, F, G; control retinas INL: 8.14 ± 0.97 and ONL: 10.5 ± 1.26 PCNA+ cells; control retinas+LPS INL: 24.71 ± 1.99 and ONL: 42.29 ± 2.38; NMDA INL: 142.53 ± 10.22 and ONL: 83.47 ± 5.80; NMDA+LPS: INL: 201.71 ± 10.129 and ONL: 112.24 ± 5.14). Because LPS increased the number of PCNA-positive cells in the NMDA-injured retinas, we investigated if this phenomenon was microglia-dependent. The transgenic *Tg(mpeg1:NTR-YFP)* line fish was treated for five days with MTZ to ablate macrophages. After two days of starting MTZ treatment, LPS was injected, and NMDA was injected three hours later. Few microglia were observed in these retinas (Figure 2E, H: NMDA ablation+LPS: 24 ± 2.48 mpeg1+ cells). PCNA-positive cell numbers were strongly reduced in the ablated and LPS treated group (Figure 2E, F, G; NMDA ablation+LPS INL: 46.43 ± 2.81 and ONL: 41.36 ± 3.24).

**Figure 2.**
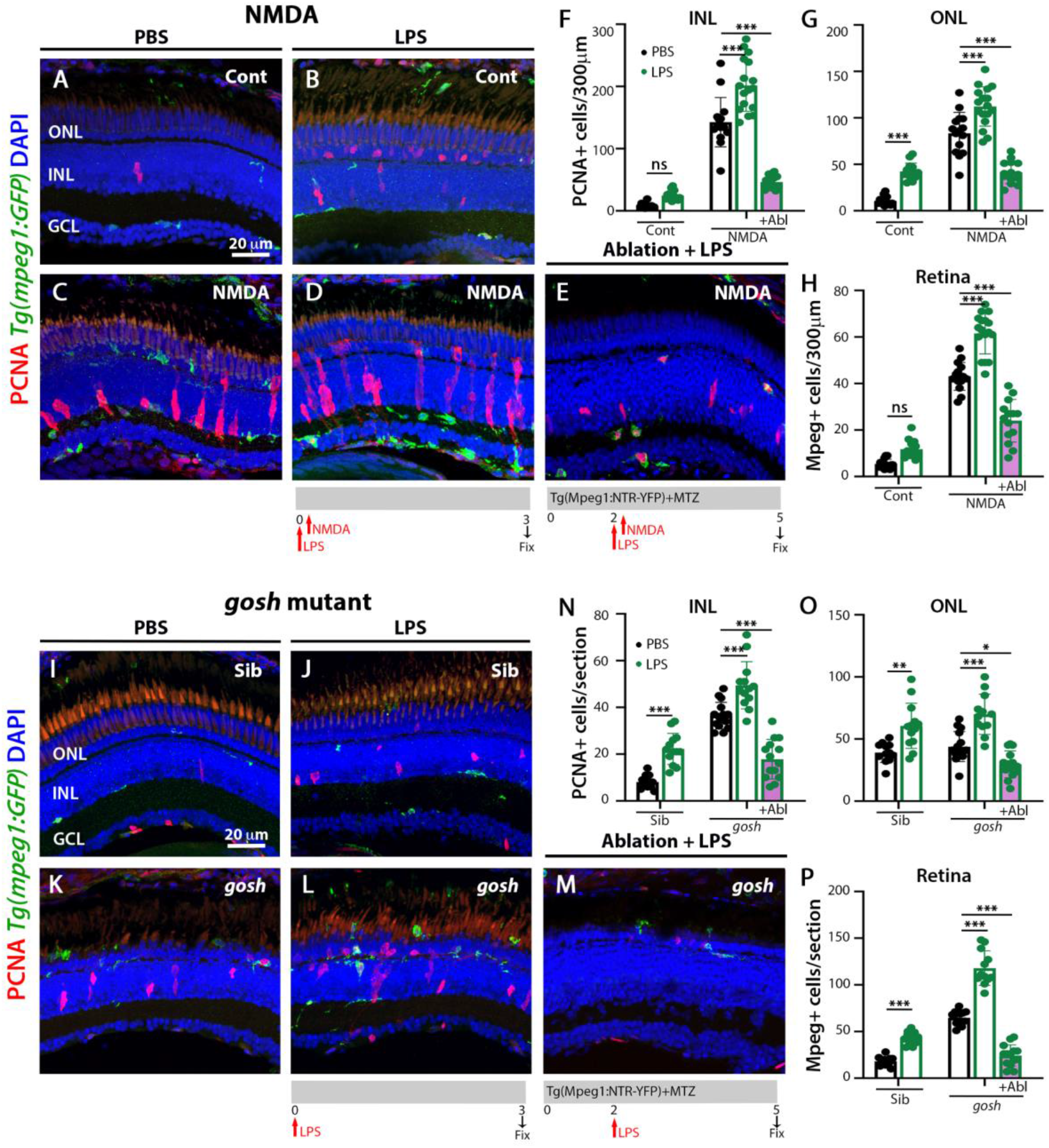
LPS-activated microglia enhance Müller glia proliferation in NMDA-injured retina and *gosh* mutant. Cryosections of 3-wpf fish with NMDA-injured at 72 hpi and *gosh* mutant retinas are labeled with an antibody against PCNA. Microglia are visualized with the *Tg(mpeg1:GFP)* transgenic line. One single injection of LPS was performed. LPS treated control samples display an increased number of cells expressing PCNA in the ONL relative to control retinas (A, B). NMDA-damaged retinas treated with LPS amplified activated microglia and proliferative cells compared to NMDA-injured retinas treated with PBS (C-D). NMDA-damaged retinas with ablated macrophages injected with LPS display few microglia and PCNA-positive cells (E). *gosh* mutant eyes injected with LPS increase microglia reactivity and PCNA-labeled cells in the INL and ONL (K, L). *gosh* mutant fish with ablated macrophages and then injected with LPS show lower number of microglia and PCNA-positive cells in the INL and ONL. Histograms display the quantification of the total number of PCNA-labeled cells (F, G) or GFP-positive macrophages (H) in 300 m of the central region of the retina for NMDA or in the entire section of the retina for *gosh* mutant (N-P). Bars and lines indicate mean ± SEM, n: 12-17. Black bars: PBS injection; green bars: LPS injection; green bars with purple fill: LPS and ablated macrophages. Two-way ANOVA with Tukey’s multiple comparisons test was applied for all the graphs (ns p>0.05; *p<0.05; **p<0.01; ***p<0.001).

Wild-type sibling and *gosh* mutant retinas also showed an increased number of inflammatory cells following LPS injection (Figure 2I-L, P; sibling retinas: 18.08 ± 1.39 mpeg1+ cells; sibling retinas+LPS 43.17 ± 1.89, p<0.001; *gosh* retinas: 64.67 ± 1.88; *gosh* retinas LPS 117.92 ± 5.32, p<0.001). Müller glia and NPCs labeled with PCNA were increased in sibling and the *gosh* mutant upon LPS treatment (Figure 2I-L, N, O, sibling retinas INL: 8.08 ± 0.82 and ONL: 39.08 ± 2.29 PCNA+ cells; sibling retinas+LPS INL: 22.23 ± 1.85, p<0.001; and ONL: 60.62 ± 5.05, p<0.001; *gosh* mutant INL: 36.47 ± 1.48 and ONL: 43.87 ± 3.10; *gosh* mutant+LPS INL: 49.47 ± 2.81, p<0.001 and ONL: 69.92 ± 4.7, p<0.001). The *gosh* mutant fish combined with the *Tg(mpeg1:NTR-YFP)* line were treated with MTZ for five days. After two days of MTZ treatment, the *gosh* mutant fish were injected with LPS (Figure 2M). The nitroreductase-MTZ system allows for efficient microglia ablation (Figure 2M, P: *gosh* mutant ablation+LPS: 24.29 ± 3.05 mpeg1+ cells). Ablated and LPS-injected retinas displayed few PCNA-positive cells (Figure 2M-O; *gosh* mutant ablation+LPS INL: 17.86 ± 2.26 and ONL: 29.64 ± 2.89). These data suggested that the inflammation induces the proliferation of Müller glia in the wild-type retina and potentiates a regenerative response in the damaged retina via the action of macrophages/microglia.

### Differential gene expression profile of pro-inflammatory and anti-inflammatory molecules in acute and chronic damage models

To uncover the inflammatory state during the regenerative response in the acute and chronic damaged retina, we assessed relative gene expression levels of inflammatory cell genes. Expression of the mpeg1 and p2ry12 genes was used to monitor microglia/macrophage cell activation, and immune genes known to induce a pro-inflammatory (il-1β, tnf-α, tnf-β) or anti-inflammatory/remodeling (il-10, tgf-β1, arg1, acr2) immune response were evaluated. RNAs from NMDA-injured retina at different times (6 hours to 1-week post-injection) were used to perform quantitative real-time PCR (qPCR) (Figure 3A-C). NMDA injection induced the activation of macrophage/microglia at 24 hpi and remained at elevated levels at 72 hpi. After 1 week, p2ry12 gene expression went down to a similar value to control, but mpeg1 gene expression remained higher than control (Figure 3A). In NMDA injected retina samples, gene expression levels of the pro-inflammatory cytokine il-1β peaked at 6 hpi (~3-fold, p<0.05) and then returned to control levels (Figure 3B). tnf-α gene expression increased from 6 hpi and tnf-β from 12 hpi, their expression peaks were observed at 24-48 hpi (~4-fold for tnf-α, p<0.001; ~5-fold for tnf-β, p<0.001). At 1 week post-injury, tnf-α and tnf-β still remained significantly higher than control levels (~1.6-fold, p<0.05 for tnf-α and tnf-β). il-10 and ccr2 genes displayed a similar pattern with two significantly elevated peaks, one at 24 hpi (~2.5-fold for il-10, p<0.01; ~3-fold for ccr2, p<0.01) and the second at 72 hpi (~2-fold for il-10, p<0.05; ~2-fold for ccr2, p<0.05). The anti-inflammatory cytokine arg1 gene was expressed at higher levels than control at 24 and 48 hpi (approximately 2-fold at 24 hpi, p<0.05, and 3-fold at 48 hpi, p<0.01), while tgf-β1 showed similar levels to control samples through all the time points evaluated. All the anti-inflammatory genes evaluated in this study were reduced to control levels by 1-week post-injury.

**Figure 3.**
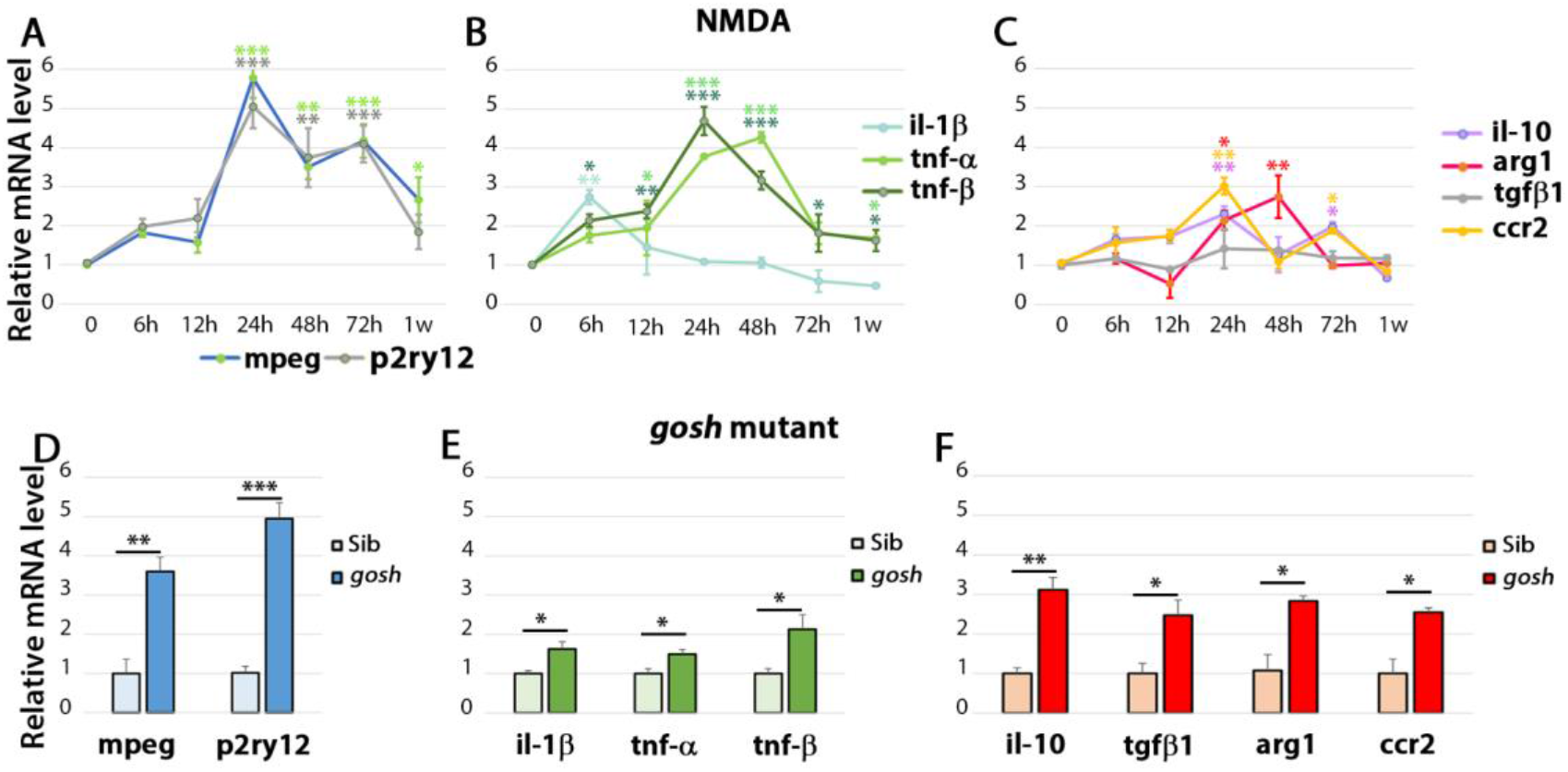
The acute injury model induces pro-inflammatory cytokine gene expressions that switch to anti-inflammatory cytokine expressions, whereas chronic injury shows the simultaneous expression of pro-inflammatory and anti-inflammatory gene expressions. Relative mRNA expressions of mpeg1 and p2ry12 (A, D); IL-1β, TNF-α, and TNF-β (B, E); IL-10, Tgf-β1, Ccr2, and Arg1 (C, F) in NMDA-damaged retina and *gosh* mutant at 3 wpf were measured by qPCR. There is significant upregulation of mpeg1 and p2ry12 in acute and chronic injured eyes. The time course of NMDA-damaged retinas depicts an early IL-1β peak, followed by stimulation of TNF-α and TNF-β gene expressions. This upregulation tendency falls gradually within 1 week. Anti-inflammatory cytokine gene expressions are induced from 24 hpi, and levels are back to control at 1 week, except for Tgf-β1 that always remains at basal levels. *gosh* mutants display a small but significant upregulation of pro-inflammatory cytokine gene expressions, whereas anti-inflammatory cytokines are upregulated. Graphs represent the mean value of 2-3 independent experiments ± SEM. For the NMDA experiments, a One-way ANOVA with Dunnett’s multiple comparisons test was employed; for *gosh* mutant results, an unpaired Student’s t-test was applied. Statistical significance between bars is indicated *p < 0.05, **p < 0.01, ***p < 0.001.

Since the *gosh* mutant is a chronic degeneration mutant, only one-time point was assessed (Figure 3D-F). Inflammatory cell markers were induced in the *gosh* mutant (~4-fold for mpeg1, p<0.01; ~5-fold for p2ry12, p<0.001). Pro-inflammatory cytokine genes were upregulated, but the fold increase was slight (~1.7-fold for il-1β and tnf-α, p<0.05; ~2-fold for tnf-β, p<0.05). Anti-inflammatory cytokines were all upregulated with a fold increase of approximately 3-fold. It is worth noting that tgf-β1 was upregulated in the chronic damage, but not in the acute damage, suggesting an expression dependent on the injury paradigm. Hence, in the acute retinal damage, the immune response is biphasic with an initial pro-inflammatory phase, followed by an anti-inflammatory/remodeling phase. In chronic retinal damage, both pro-inflammatory and anti-inflammatory cytokines overlap, with pro-inflammatory cytokines showing a slight increase, while pro-inflammatory showing a larger fold increase.

### Microglia express TNF-α in acute and chronic retinal damage in zebrafish

A reporter transgene for TNF-α, which is a pro-inflammatory cytokine and a well-established marker of pro-inflammatory macrophages, was demonstrated to be helpful in discriminating macrophage subsets in zebrafish (Nguyen-Chi et al., 2015). To identify pro-inflammatory macrophages, we used the *TgBAC(tnfα:GFP)* and *Tg(mfap4:tdTomato-CAAX)* double-transgenic reporter lines and injected NMDA into the eye (Figure 4A-J”) or introduced then into the *gosh* mutant background (Figure 4K-N”). NMDA-damaged retinas were evaluated at 0, 12, 24, 48, and 96 hours following NMDA injection. Control retinas displayed microglia expressing tdTomato and their cell shape was thin and ramified, TNF-α drove GFP expression in amacrine cells based on their localization and shape, but was not detected in microglia (Figure 4A-B”). At 12 hours after NMDA injection, microglia increased their numbers and appeared to be activated, by 24 hpi, the recruitment and cell shape of microglia showed strong reactivity (Figure 4C-D”). After 12 and 24 hpi, microglia expressed GFP and tdTomato, with variation in the expression. Some tdTomato-positive microglia/macrophages expressed high or medium levels of TNF-α, while others did not express TNF-α, and few microglia expressed only TNF-α. At 48 hpi, TNF-α pro-inflammatory microglia/macrophages were still detected in the retina. By 96 hpi, microglia mfap4-expressing cells did not express tnf-α:GFP, however, these microglia showed reactivity features similar to the wild-type retinas (Figure 4I-J”). The *gosh* mutant presented a high density of amoeboid-shaped microglia in the degenerating photoreceptor layer and outer segments relative to wild-type sibling retinas (Figure 4K-N”). Few *gosh* mutant microglia co-labeled with tnf-α:GFP. Thus, NMDA-injured retinas displayed a population of pro-inflammatory microglia/macrophage from 12 to 48 hpi, which were no longer detected at 96 hpi. These expression profiles might represent the switch of pro-inflammatory to anti-inflammatory/resolution of microglia/macrophage. On the other hand, in the *gosh* mutant, microglia/macrophage showed an activated state, with some cells expressing the pro-inflammatory cytokine TNF-α.

**Figure 4.**
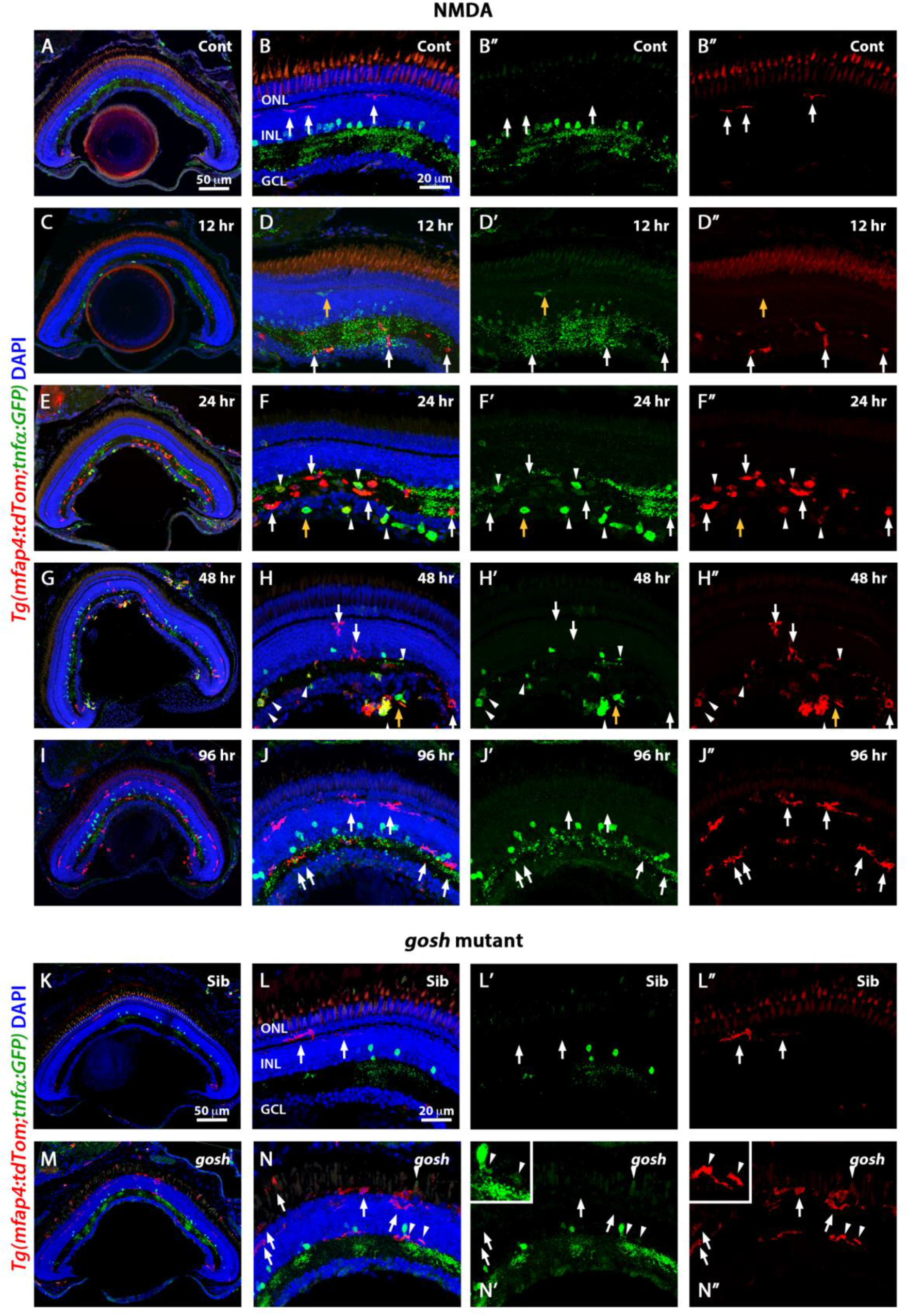
M1-like pro-inflammatory microglia are transiently distinguished in acute injury and low levels in chronic injury retina models. Double-transgenic fish *Tg(mfap4:tdTomato;tnfα:GFP)* were used to monitor M1-like pro-inflammatory microglia. Confocal images show that the control sibling presents tdTomato expression, while GFP appears to be expressed by amacrine cells (A, B”). NMDA-injured retinas were evaluated at 12, 24, 48, and 96 hours (C-J). After 6 hours of NMDA injection, retinas increase microglia density, and after 12 hours, microglia start to co-express TNF-α. TNF-α macrophages are detectable up to 72 hpi, and they are no longer detectable by 96 hpi. Wild-type sibling retinas display few and ramified microglia (K, L”). *gosh* retinas exhibit many ameboid microglia in the ONL and outer segments layers, and few of these microglia express TNF-α (M, N”). White arrows: tdTomato-positive microglia; yellow arrows: TNF-α-positive microglia; arrowhead: double-positive tdTomato;TNF-α.

### Modulating inflammation modifies microglia populations in acute and chronic retinal models

NMDA-injured and *gosh* mutant retinas treated with the anti-inflammatory drug dexamethasone displayed a low number of proliferating Müller glia (Figure 1). On the contrary, this inflammatory signal LPS stimulated the number of proliferating Müller glia (Figure 2). To assess whether dexamethasone and LPS affect the pro-inflammatory microglia/macrophage phenotype in the injured retina, we used the transgenic reporter lines *TgBAC(tnfα:GFP)* and *Tg(mfap4:tdTom)*. Control retinas showed microglia, which did not express TNF-α (Figure 5A). In contrast. NMDA-induced retinal damage displayed pro-inflammatory microglia (Figure 5B). Dexamethasone treatment resulted in microglia that did not co-label with GFP (Figure 5C, D). LPS treatment induced pro-inflammatory microglia in the undamaged retina, while the NMDA-injured retina showed tdTomato and GFP co-expression in the microglia (Figure 5E, F). Sibling and *gosh* mutant retinas possessed microglia in the retina (Figure 5G, H). Upon dexamethasone treatment, sibling and *gosh* retinas revealed microglia that did not express tnfα:GFP (Figure 5I, J). LPS treatment stimulated microglia to express GFP in sibling and *gosh* retinas (Figure 5K, L). Taken together, these data suggested that dexamethasone inhibited pro-inflammatory microglia/macrophage in acute and chronic retinal damage, whereas LPS induced pro-inflammatory microglia in acute and chronic injured retinas. Thus, dexamethasone reduced the proliferative Müller response, which did not have M1-like microglia. On the contrary, LPS stimulated the proliferative Müller response, accompanied by M1-like microglia in acute and chronic injured retina models.

**Figure 5.**
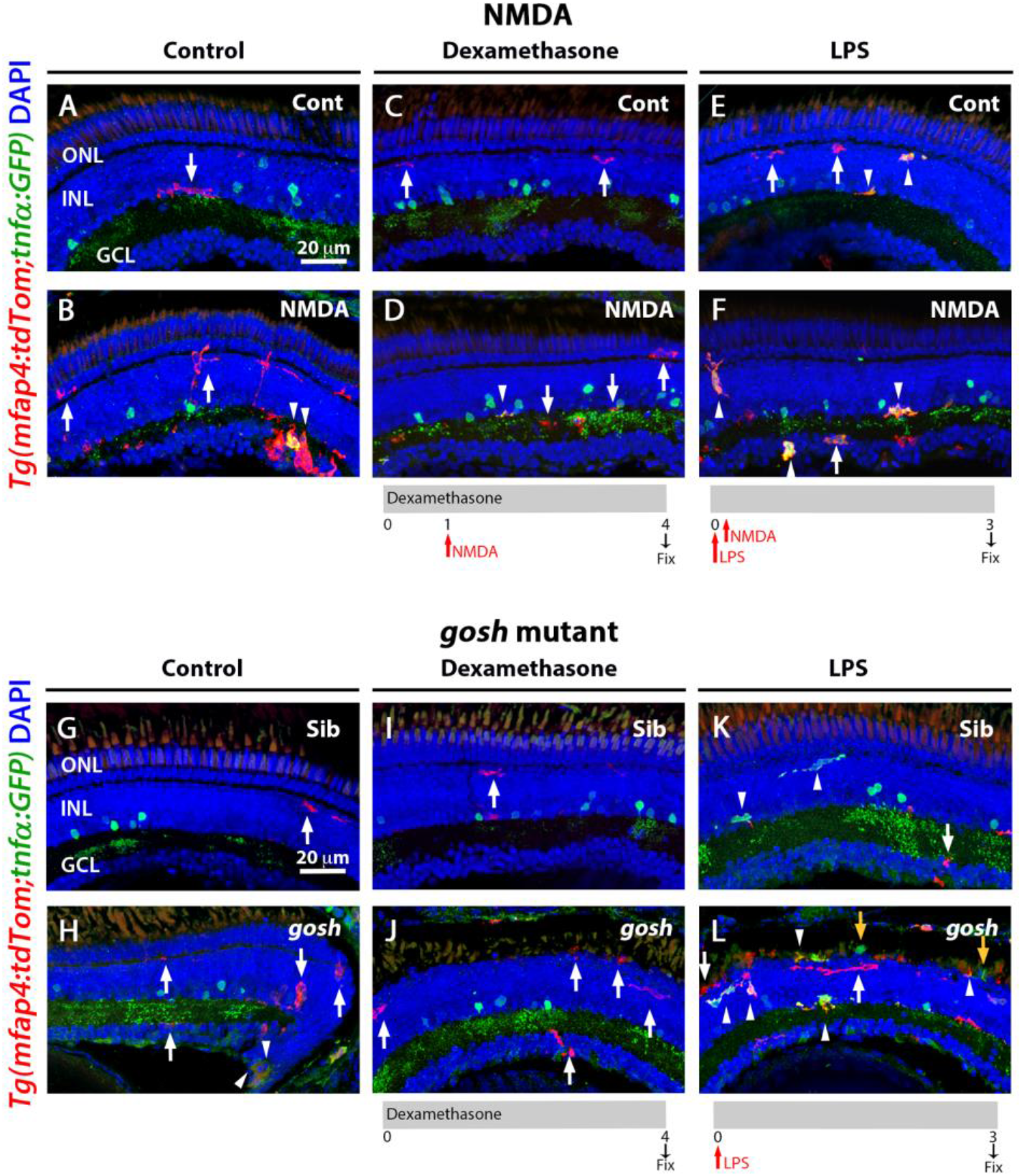
M1-like microglia are modulated by dexamethasone and LPS treatment in acute and chronic damaged retinas. Double-transgenic fish *Tg(mfap4:tdTomato;tnfα:GFP)* allows the visualization of M1-like pro-inflammatory microglia in retinas treated with dexamethasone or LPS. Control retinas show some ramified tdTomato-positive microglia (A, arrow). NMDA-injured retinas show several ameboid microglia, and some express TNF-α (B, arrowhead). Dexamethasone treatment reduces microglia accumulation and ameboid shape. Few microglia co-express GFP and tdTomato (C, D). LPS treatment increases the presence of microglia, including expression of TNF-α in control (E) and NMDA-damaged retinas (F). *gosh* mutants display ameboid tdTomato microglia, and some also express TNF-α:GFP (H). Dexamethasone treatment prevents TNF-α expression in sibling (I) and *gosh* mutant (J). LPS injection increases tdTomato-positive microglia and TNF-α/tdTomato-double-positive cells in control (K) and *gosh* mutant (L). White arrows: tdTomato-positive microglia; yellow arrows: TNF-α-positive microglia; arrowhead: double-positive tdTomato;TNF-α.

## Discussion

We investigated the role of inflammation in retina regeneration following acute or chronic injury in 3 wpf zebrafish. We showed that these different injury paradigms induced inflammation and proliferation of Müller glia and NPCs. Modulating inflammation regulated the proliferative response. Acute retinal damage induced an early upregulation of pro-inflammatory cytokines and later increased expression of anti-inflammatory cytokines. In addition, TNF-α expressing M1-like pro-inflammatory microglia/macrophage were transiently present during the regenerative response. On the other hand, chronic retinal damage upregulated a robust anti-inflammatory response in the presence of a mild pro-inflammatory response. TNF-α expressing M1-like microglia are modulated by dexamethasone and LPS treatment in both the acute and chronic injury paradigms, which affected Müller glia proliferation (Figure 6).

**Figure 6.**
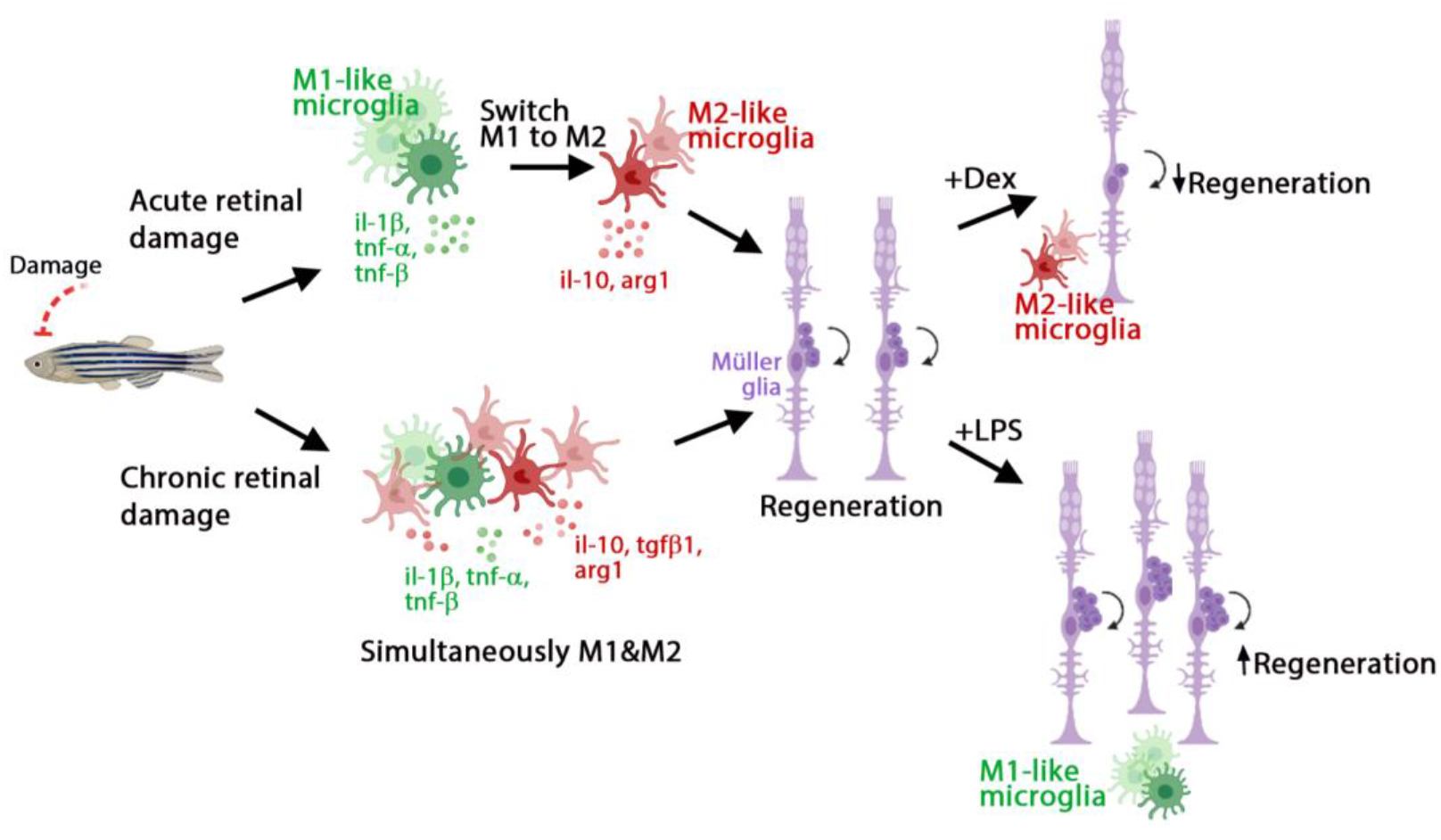
Model of inflammation regulating the regenerative response in acute and chronic retinal damage in zebrafish. Diagram representing microglia/macrophage activation and polarization upon retinal damage in zebrafish to induce efficient regeneration. Acute injury stimulates an early pro-inflammatory response, which then switches to an anti-inflammatory profile. Chronic injury stimulates a dominant anti-inflammatory response, accompanied by a mild pro-inflammatory response. Modulating inflammation can inhibit or amplify the regeneration capacity via regulating the presence of M1-like pro-inflammatory microglia.

Tissue injury elicits a rapid and strong immune response in zebrafish, which is required for regeneration (Baumgart, Barbosa, Bally-Cuif, Gotz, & Ninkovic, 2012; Iribarne, 2021; Lai, Marin-Juez, & Stainier, 2019; Marz, Schmidt, Rastegar, & Strahle, 2010; Nagashima & Hitchcock, 2021). Recent research had shown that the knockdown of macrophage differentiation impaired fin regeneration in larvae, and genetic ablation of macrophages affected fin growth in adult zebrafish (Li, Yan, Shi, Zhang, & Wen, 2012; Petrie et al., 2014). Blocking inflammation in spinal-lesioned zebrafish larvae reduced axonal bridging and GFAP-positive proliferating radial glia (Ohnmacht et al., 2016; Tsarouchas et al., 2018), whereas activation of the immune system increased axonal regeneration (Tsarouchas et al., 2018). Depleting microglia or inhibiting their functions in the damaged larval and adult zebrafish retina blocked the Müller glia regenerative response (Silva et al., 2020; White et al., 2017; Zhang et al., 2020). Our results are consistent with the literature, as the injury paradigms used in this research induced damage that triggered microglia/macrophage reactivity. Moreover, inhibiting the inflammatory response or ablating macrophages reduced the number of proliferating Müller glia and NPCs, while activating the inflammatory response increased the number of proliferating Müller glia and NPCs in both acute and chronic retinal injury models. These results suggest that macrophages play a critical role in zebrafish tissue regeneration. Macrophages exist in a variety of states that are acquired under different environments. In our acute damage model, we observed that TNF-α expressing M1-like pro-inflammatory macrophages transiently accumulated in the early regenerating retina, whereas later, M2-like anti-inflammatory macrophages were detected. As expected, expression of pro-inflammatory cytokine genes was upregulated early, followed by the upregulation of anti-inflammatory cytokines. In zebrafish, a limited number of studies described the presence of different macrophage subtypes during the regeneration process. Petrie and colleagues showed that macrophages at different stages are responsible for different functions during fin regeneration (Petrie et al., 2014). Early and transient recruitment of TNF-α expressing pro-inflammatory macrophages occurred during fin regeneration (Nguyen-Chi et al., 2017). More recent studies identified a transient pro-regenerative macrophage subtype in zebrafish with a specific gene expression profile (Cavone et al., 2021; Ratnayake et al., 2021; Sanz-Morejon et al., 2019). In the regeneration of the larval spinal cord, the pro-regenerative macrophage population expressed TNF-α (Cavone et al., 2021), however the pro-regenerative macrophage population was Wilms-positive and TNF-α-negative in heart and fin regeneration models (Sanz-Morejon et al., 2019). In contrast, a NAMP-positive pro-regenerative macrophage population was identified during muscle regeneration (Ratnayake et al., 2021). At least the TNF-α and Wilms pro-regenerative macrophage subtypes were different populations, suggesting that specific markers and macrophages subtypes are associated with regenerating different tissues. These results indicate that an inflammatory response followed by resolution is required to achieve successful regeneration, involving different macrophage subtypes in each stage. It has yet to be elucidated if there is only one macrophage subtype culpable for the scope of functions during inflammation and if there is a switch from pro-inflammatory to an anti-inflammatory profile, as was observed *in vivo* in zebrafish larvae (Nguyen-Chi et al., 2015), or whether several macrophage subtypes are involved.

Mammals undergoing chronic degeneration suffer not only primary damage by the injury, but also prolonged inflammatory activation induces secondary damage, which could be more detrimental than the primary injury itself. It has been hypothesized that inflammatory macrophages or microglia damage the CNS, while a resolving phenotype contributes to neuroregeneration (Kigerl et al., 2009). Upon traumatic spinal cord injury in rats, the M1-like macrophage response was rapidly induced and then maintained at damage sites. This response overwhelmed a comparatively smaller and transient M2 macrophage response. Interestingly, zebrafish suffering photoreceptor degeneration exhibited chronic inflammation, which had at least two components; a mild pro-inflammatory component and a stronger anti-inflammatory component. The specific cytokine environment could explain this surprising success in regeneration during chronic damage in zebrafish. The ratio between M1/M2-like macrophages could have significant implications for CNS repair. Specifically, in zebrafish, the necessity of M1-like pro-inflammatory microglia is apparent by the inhibition of this population preventing Müller glia proliferation. Alternatively, it could be that the transient pro-regenerative TNF-α expressing microglia subtype is sufficient to induce regeneration, similar to the TNF-α pro-regenerative microglia subtype stimulating spinal cord regeneration (Cavone et al., 2021). To our knowledge, our study is the first that focused on the role of microglia in modulating the regenerative response of a chronic injury in zebrafish.

A transient inflammatory response mediated by IL-1β is required for proper regeneration in the zebrafish fin fold (Hasegawa et al., 2017). Macrophages attenuated IL-1β expression and inflammation to support the survival of regenerative cells. Furthermore, inflammation mediated by IL-1β was necessary for normal fin regeneration by triggering the expression of regeneration-induced genes (Hasegawa et al., 2017). In the resected spinal cord, early expression of IL-1β promoted axonal regeneration, while prolonged high levels of IL-1β were detrimental. Another study showed that TNF-α was one of the critical signals transiently expressed by polarized macrophages during the early phases of fin regeneration (Nguyen-Chi et al., 2017). The proliferation of stromal cells depended on TNF-α/TNF-r1 signaling, suggesting that IL-1β and TNF-α are expressed at different times, with IL-1β being expressed earlier and followed by pro-inflammatory macrophages expressing TNF-α. We found that acute retinal injury induced the expression of an early peak of IL-1β, followed by increased expression of TNF-α and TNF-β during retinal regeneration. Although it remains unclear if a similar mechanism is present in both the fin and spinal cord tissues, these results suggest that the activation and duration of pro-inflammatory signals and the subsequent resolution are critical in creating an instructive microenvironment for tissue regeneration.

In conclusion, we found that acute and chronic damage stimulated a regenerative response via inducing different pro-inflammatory response strategies. Acute damage induced an early and transient pro-inflammatory response, whereas chronic damage stimulated a persistent mild pro-inflammatory response in the presence of a stronger anti-inflammatory response. Additionally, pro-inflammatory microglia/macrophages are required for the regenerative response, as their abolition impaired regeneration. Understanding how inflammation regulates regeneration in the injured zebrafish retina would provide important insights to improve the therapeutic strategies for repairing injured mammalian tissues lacking an inherent regenerative capacity.

## Material and methods

### Fish

According to standard procedures (Westerfield, 1993), zebrafish (*Danio rerio*) were maintained in the Center for Zebrafish Research at the University of Notre Dame Freimann Life Science Center. AB wild-type fish was used as a wild-type strain. The *gold rush* (*gosh)* mutant was originally isolated while screening for zebrafish visual mutants using a chemical mutagen, N-ethyl-N-nitrosourea (ENU) (Muto et al., 2005). Zebrafish transgenic lines *Tg(mpeg1:GFP)*^*gl22*^ or *Tg(mfap4:tdTomato-CAAX)*^*xt6*^ were used to monitoring microglia behavior (Ellett et al., 2011; Walton, Cronan, Beerman, & Tobin, 2015). *Tg(gfap:EGFP)*^*nt11*^ was used to visualize Müller glia (Kassen et al., 2007), *TgBAC(tnfα:GFP)*^*pd1028*^ (Marjoram et al., 2015) was used to identify pro-inflammatory macrophages, and *Tg(mpeg1:NTR-eYFP)*^*w202*^ was used to ablate macrophages (Petrie et al., 2014). Fish were euthanized by an anesthetic overdose of 0.2% 2-phenoxyethanol, and eyes were enucleated for further processing. All experimental protocols were approved by the animal use committee at the University of Notre Dame and followed the National Institutes of Health guide for the care and use of Laboratory animals (NIH Publications No. 8023, revised 1978).

### Optokinetic response (OKR) behavior test

*gosh* visual mutants, obtained from incrossing *gosh* carriers, were identified at 5-7 days-post-fertilization (dpf), using the OKR test (Iribarne et al., 2017). In a petri dish containing methylcellulose, 10 wells were filled with aquarium water in which the fish were raised to minimize stress to the fish. Individual larvae were located into single wells to be partially immobilized and allow examination under a stereoscopic microscope. To evaluate visual acuity, a drum with black and white vertical stripes (at 18° separation) was placed around the petri dish and spun at 10-20 rpm. Larval eye movement was observed under the stereoscopic microscope to identify cone blind fish.

### NMDA lesion

Retinal lesions were induced by a single intravitreal injection of 0.5 - 1 nL of freshly prepared 100 mM N-Methyl-D-aspartic acid (NMDA, M3262, Sigma). Briefly, fish were anesthetized in 0.1% 2-phenoxyethanol, and under microscopic visualization, NMDA was delivered using a FemtoJet express microinjector (Eppendorf).

### Drug treatment

Dexamethasone (Dex, D1756, Sigma-Aldrich) was diluted in 100% methanol to generate a 5 mM stock concentration. The 5 μM working concentration was prepared by diluting the stock solution in aquarium water. Fish were placed in static tanks, and the solution was changed once a day. Lipopolysaccharides from Escherichia coli O55:B5 (LPS, L2880, Sigma-Aldrich) were dissolved in PBS to a concentration of 1 mg/mL. One single intravitreal injection was carried out using a FemtoJet express microinjector (Eppendorf).

### Ablation of macrophages

To ablate macrophages, the transgenic line *Tg(mpeg1:NTR-eYFP)*^*w202*^ was used (Petrie et al., 2014). 5 mM Metronidazole (MTZ, M3764, Sigma-Aldrich) was prepared fresh and covered from light. 3-wpf fish were housed in static tanks with fish water containing MTZ, and the solution was changed every day.

### Histology

Head samples were fixed with 4% paraformaldehyde in 0.1 mol/L phosphate buffer (4% PFA, Sigma-Aldrich), pH 7.3 overnight at 4°C, or in 9:1 ethanol:formaldehyde (Fisher Scientific) for 2 hours at room temperature in a shaker. Specimens were washed in PBS three times (4% PFA), or previously washed with a gradient of ethanol and then PBS for 9:1 ethanol:formaldehyde, then cryoprotected and rapidly frozen. Immunolabeling of cryosections (14μm thickness) was performed as previously described (Masai et al., 2003). For antigen retrieval, cryosections were pretreated with heat (~90°C, 10 minutes, in 10 mM citrate buffer pH 6.0). Mouse anti-PCNA antibody (clone PC10, Sigma P8825; 1:1000) and mouse anti-4C4 (HPC Cell Cultures, 92092321, 1:50) were used. GFP (Life Technology, A11122, 1:200) and RFP (Rockland, 600-401-379, 1:200) antibodies were used to amplify the signal or detect GFP and tdTomato-CAAX after antigen retrieval. Nuclear staining was performed using 5μg/mL DAPI (Invitrogen). Images were scanned using a confocal laser scanning microscope (Nikon A1r).

### Quantitative real-time PCR

RNA was isolated from 3-wpf NMDA, *gosh* mutants, and controls fish. 12-15 fish heads were dissected and pooled. RNA was extracted using TRIzol reagent (Life Technologies). Total cDNA was synthesized from 1 μg of RNA using qScript cDNA SuperMix (Quanta Biosciences, Gaithersburg, MD). Reactions were assembled using PerfeCta SYBR Green SuperMix (ROX; Quanta Biosciences). Primers used in this study are included in the following table. Data were acquired using the StepOnePlus Real-Time PCR system (Applied Biosystems, Foster City, CA). Analysis was performed using the Livak 2^−ΔΔC(t)^ method.

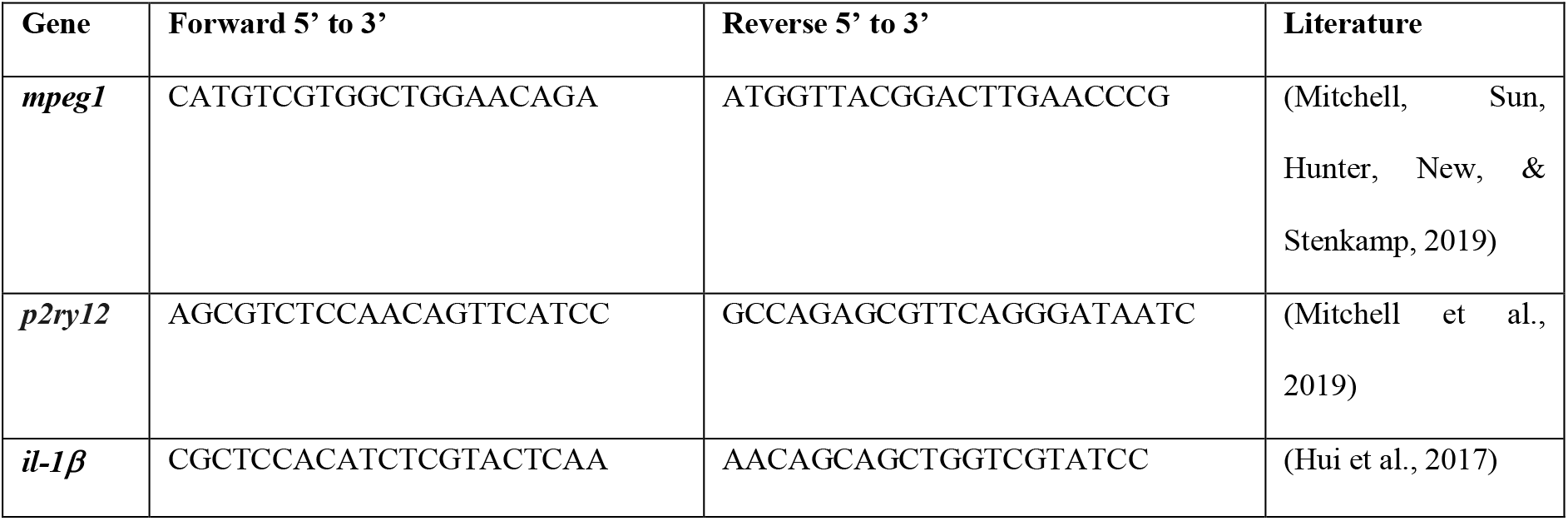
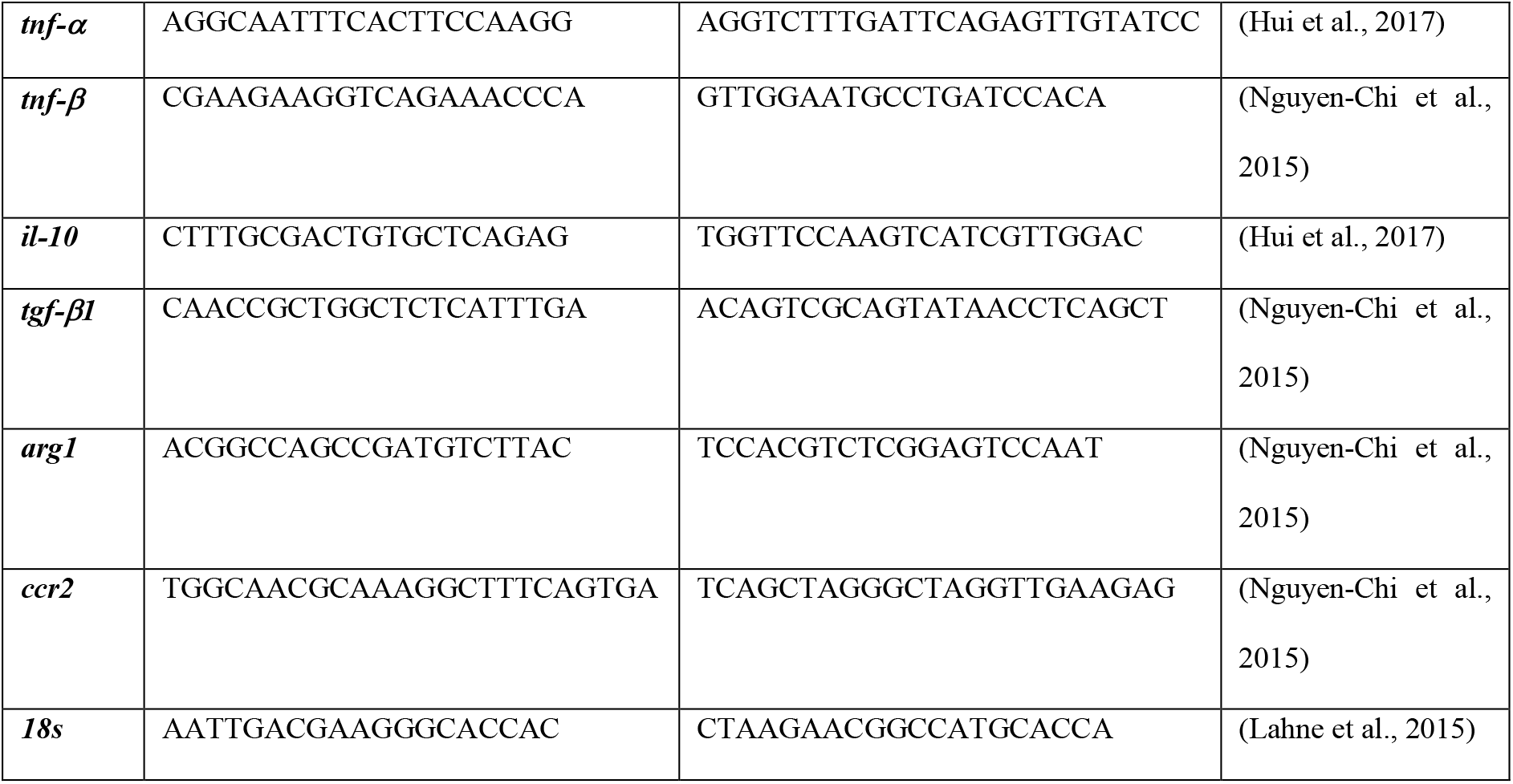

### Quantifying cells

To quantify the PCNA-labeled cells, we count the number of cells in the outer nuclear layer (ONL) and the inner nuclear layer (INL), or mpeg1-expressing cells in all the retinal layers throughout the depth of the z-stack in a line of 300 μm length of the NMDA-damaged retinas or from an entire retinal section for *gosh* mutant. A total of 3-7 fish were used for treatment, and every experiment was performed at least three times. Counts were then averaged and the SEM was calculated. The statistical significance of differences of single comparisons between the control and the treatment group was evaluated with a two-tailed, unpaired Student’s t-test (Figure 3D-F) and one-way ANOVA with Dunnett’s multiple comparison test to compare all the values against the control value (Figure 3A-C). The differences between multiple groups were determined using one-way ANOVA with Tukey’s *post-hoc* test (Figure 1) and two-way ANOVA (Figure 2). The graphs were performed using Graphpad Prism9 software and Excel.

## Acknowledgements

Zebrafish transgenic lines *Tg(mpeg1:GFP)* and *Tg(mfap4:tdTomato-CAAX)* were a generous gift from Nishtha Ranawat and Ichiro Masai, *Tg(mpeg1:NRT-eYFP)* were kindly provided by Anne Huttenlocher and Sofia de Oliveira.

---

**Supplementary Figure 1.**
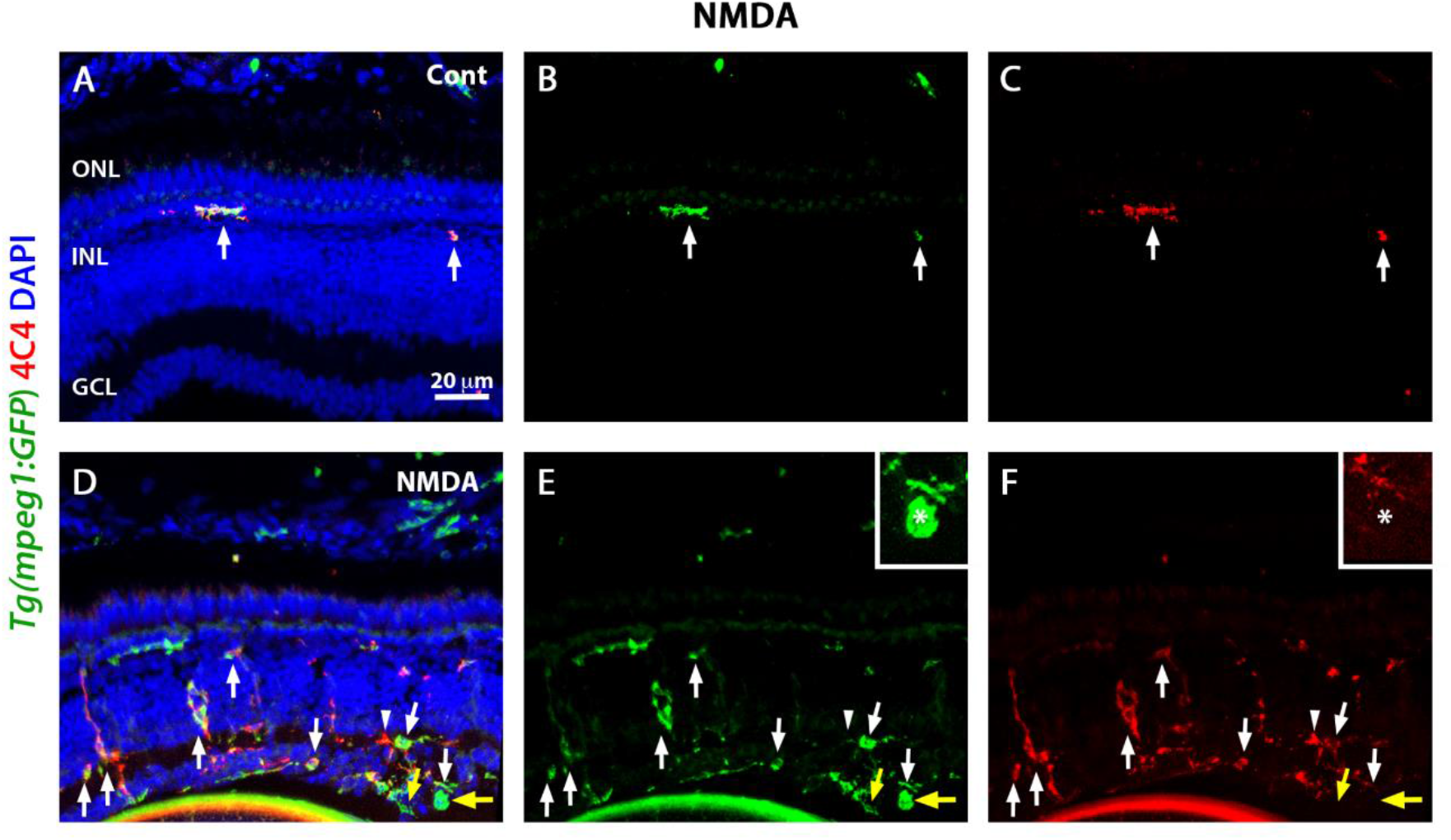
NMDA damage recruits microglia and few peripheral macrophages. Seventy-two hours after injection, control and NMDA-injured retinas expressing the transgene *Tg(mpeg1:eGFP*), which visualizes macrophages, were stained with the antibody against 4C4 to label microglia, but not peripheral macrophages. Nuclei were labeled with DAPI (blue). In control retinas, GFP and 4C4 label the same population of cells (A-C, arrows). NMDA-injured retinas display most GFP-positive cells that are co-label with 4C4 (D-F, arrows). Few cells are infiltrated peripheral macrophages (yellow arrows, inset). Arrow: 4C4- and GFP-double positive cells; yellow arrow: GFP-positive cells (shown in the inset); arrowhead: 4C4-positive cells; asterisks in the inset: GFP-positive cells.

**Supplementary Figure 2.**
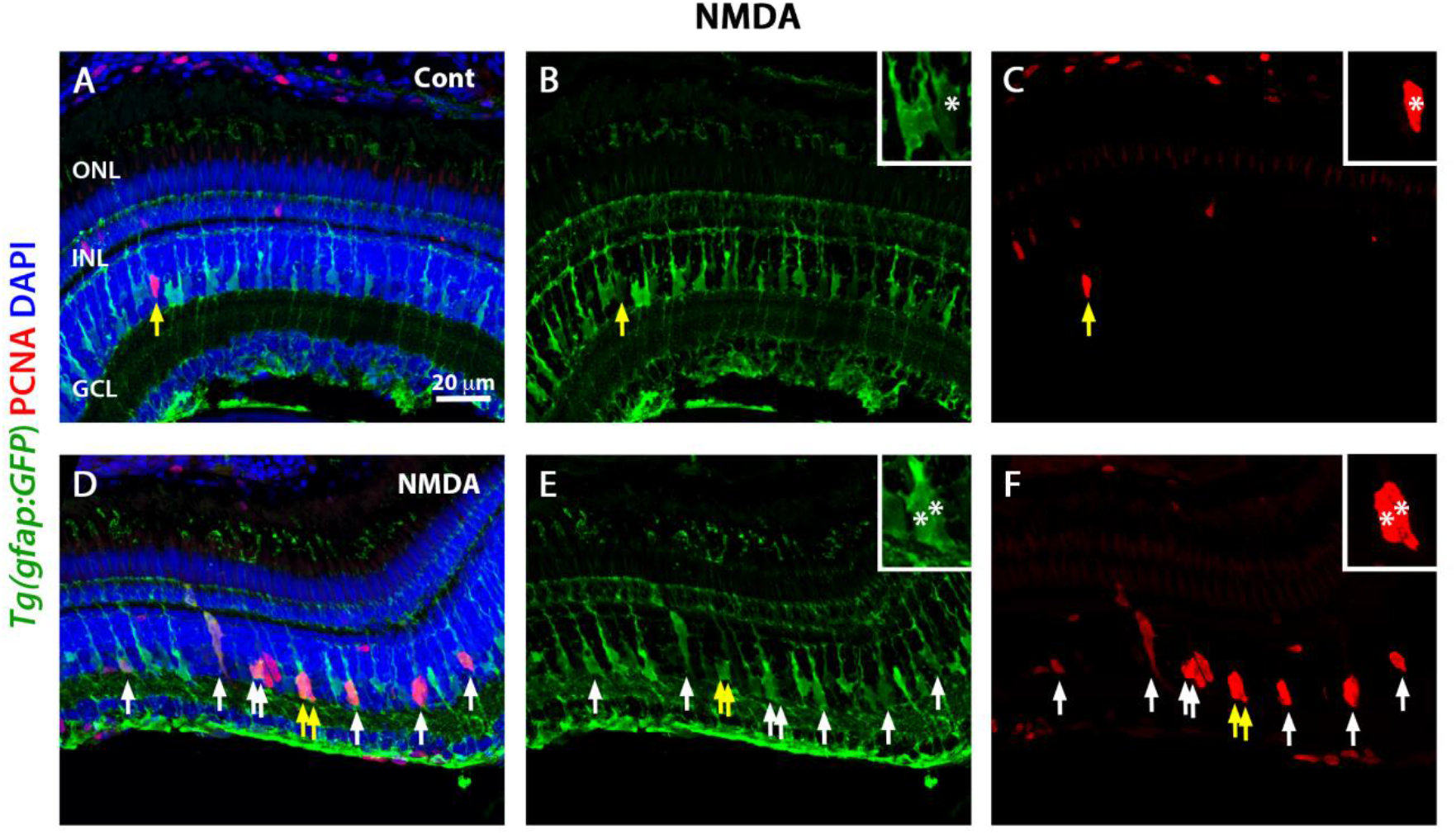
Müller glia proliferate in the acute retinal damage in zebrafish. Forty-eight hours after injection, control and NMDA-injured retinas expressing the transgene *Tg(gfap:eGFP*), which visualizes Müller glia, were stained with the antibody against PCNA. Nuclei were counterstained with DAPI (blue). In control retinas, GFP expression is observed in Müller glia cell bodies and apical-basal extended processes (A-C). Some Müller glia and cells in the ONL express PCNA indicating persistent neurogenesis (arrow). NMDA-injured retinas display several GFP-positive cells labeled with PCNA (D-F, arrows). Arrow: PCNA- and GFP-double positive cells; yellow arrow: cells shown in the inset; asterisks in the inset: PCNA- and GFP-double positive cells.

**Supplementary Figure 3.**
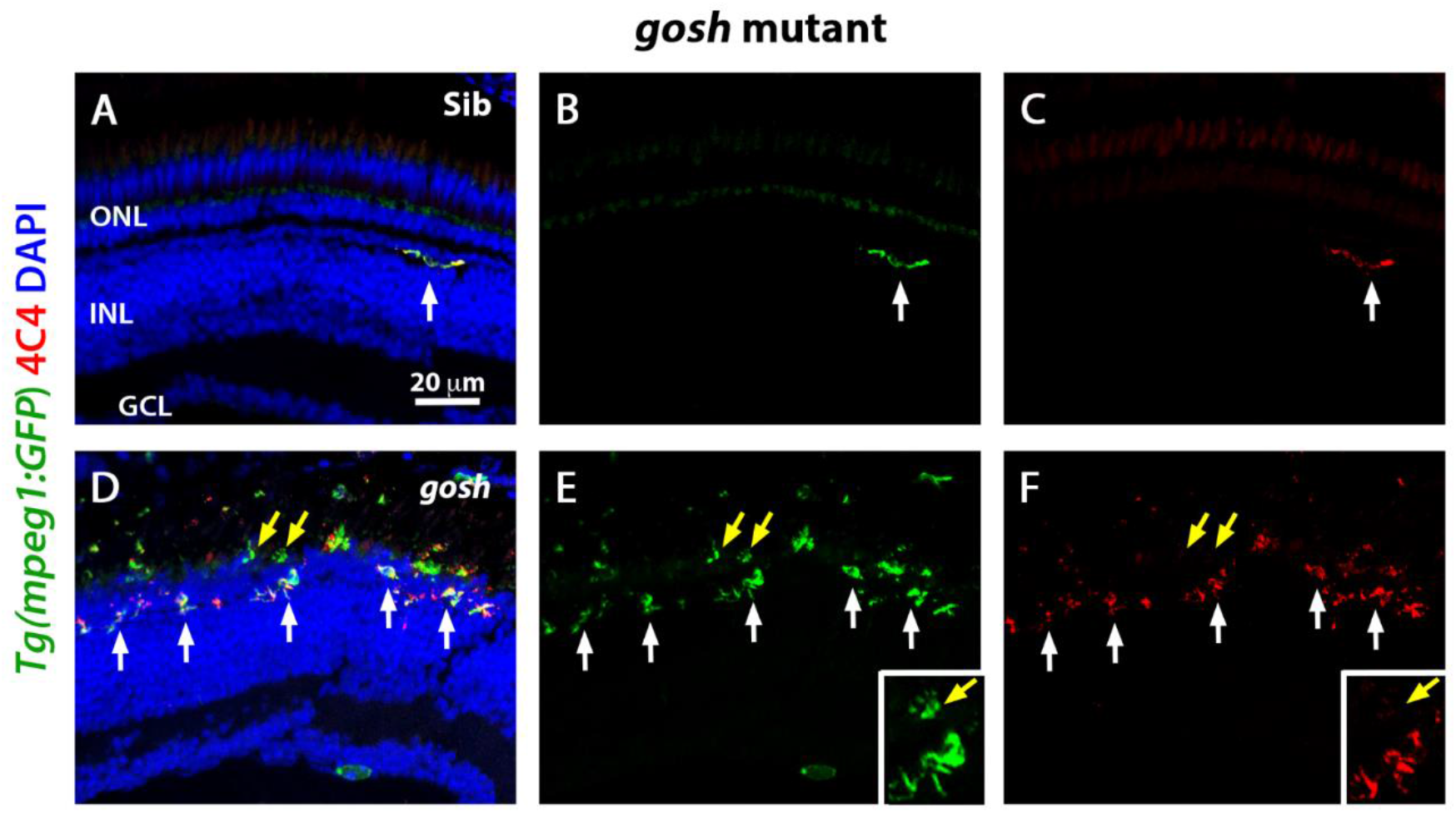
Microglia and few peripheral macrophages are detected in the *gosh* mutant. Three-wpf wild-type sibling and *gosh* mutant retinas combined with the transgene *Tg(mpeg1:eGFP*) were stained with the antibody against 4C4. Nuclei were marked with DAPI (blue). In wild-type retinas, GFP and 4C4 labels are observed in microglia (A-C, arrow). *gosh* mutants show GFP-positive cells stained with 4C4 (D-F, arrows). Few cells are stained only with GFP, which indicates that they are peripheral macrophages (yellow arrows, inset). Arrow: 4C4- and GFP-double positive cells; yellow arrow: GFP-positive cells (shown in the inset); arrowhead: 4C4-positive cells; asterisks in the inset: GFP-positive cells.

**Supplementary Figure 4.**
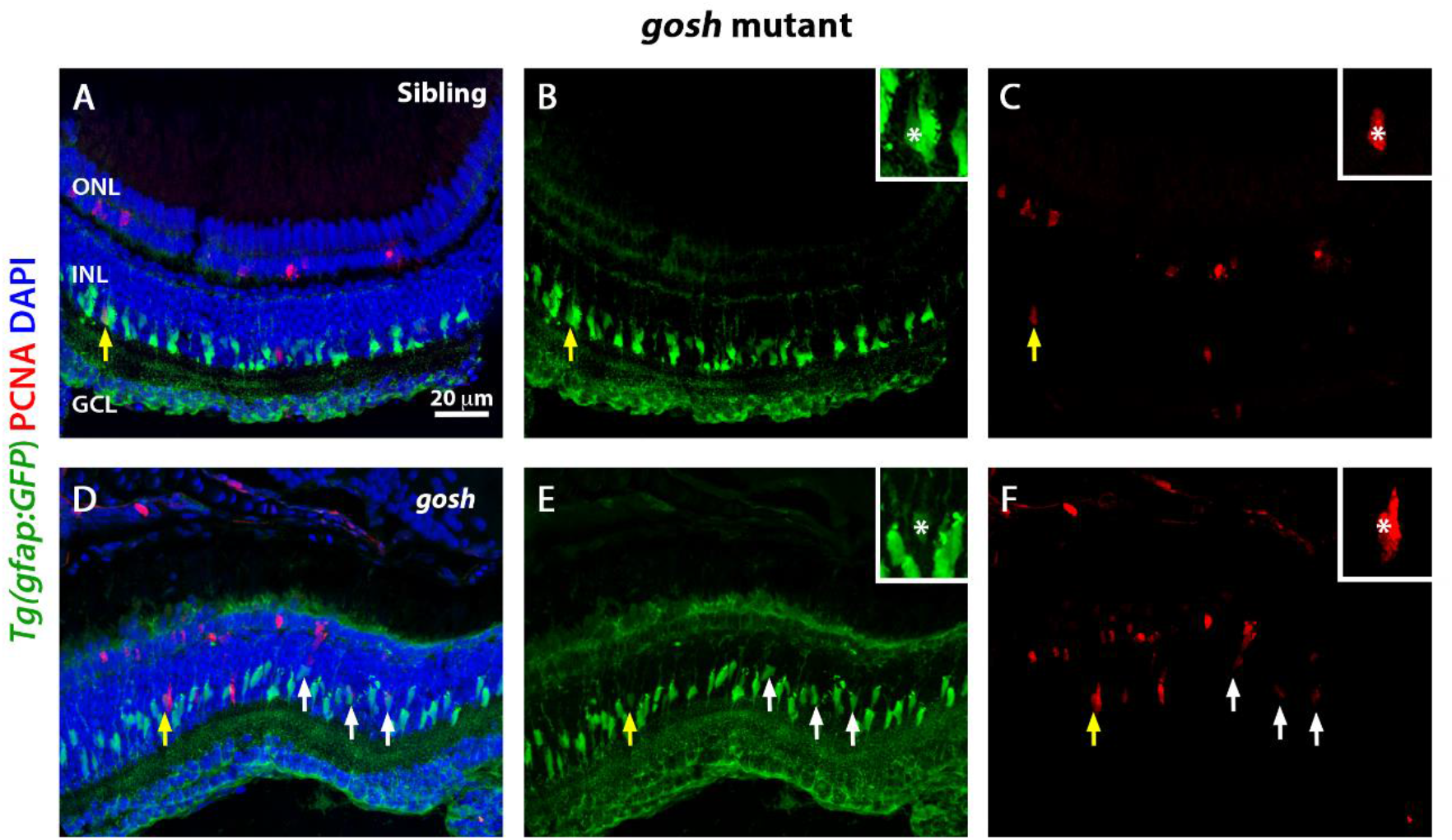
Müller glia proliferate in the chronic retinal damage in zebrafish. Three-wpf wild-type sibling and *gosh* mutant retinas combined with the transgene *Tg(gfap:eGFP*), which visualizes Müller glia, were stained with the antibody against PCNA. Nuclei were counterstained with DAPI (blue). In wild-type retinas, GFP expression is observed in Müller glia cell bodies and processes (A-C). Some Müller glia and cells in the ONL express PCNA indicating persistent neurogenesis. *gosh* mutants show several GFP-positive cells stained with PCNA (D-F, arrows). Arrow: PCNA and GFP-double positive cells; yellow arrow: cells shown in the inset; asterisks in the inset: PCNA- and GFP-double positive cells.

